# Gut Microbiome Functional and Taxonomic Diversity within an Amazonian semi-nomadic hunter-gatherer group

**DOI:** 10.1101/590034

**Authors:** LC Conteville, J Oliveira-Ferreira, AC Vicente

**Affiliations:** Laboratory of Molecular Genetics of Microrganisms, Oswaldo Cruz Institute/Fiocruz, Av. Brasil 4365, Manguinhos, Rio de Janeiro, Brazil; Laboratory of Immunoparasitology, Oswaldo Cruz Institute/Fiocruz, Av. Brasil 4365, Manguinhos, Rio de Janeiro, Brazil

**Keywords:** Gut microbiome, hunter-gatherers, semi-nomadic, Yanomami, Amerindian, westernization, biomarkers, functionality, taxonomic

## Abstract

**Background:** Human gut microbiome profiles have been associated with human health and disease. These profiles have been defined based on microbes’ taxonomy and more recently, on their functionality. Human groups that still maintain traditional modes of subsistence (hunter-gatherers and rural agriculturalists) represent the groups non-impacted by urban-industrialized lifestyles, and therefore study them provide the basis for understanding the human microbiome evolution. The Yanomami is the largest semi-nomadic hunter-gatherer group of the Americas, exploring different niches of the Amazon rainforest in Brazil and Venezuela. In order to extend the analysis of this unique and diverse group, we focused on the gut microbiome of the Yanomami from Brazil and compared with those from Venezuela, and also with other traditional groups from the Amazon, considering taxonomic and functional profiles.

**Results:** A diversity of taxonomic biomarkers were identified to each South American traditional group studied, including the two Yanomami groups, despite their overall similarity in the taxonomic gut microbiome profiles. Broader levels of functional categories poorly discriminated traditional and urban-industrialized groups. Interestingly, a diversity was observed with the stratification of these categories, clearly segregating those groups. The Yanomami/Brazil gut microbiome presented unique functional features, such as a higher abundance of gene families involved in regulation/cell signaling, motility/chemotaxis, and virulence, contrasting with the microbiomes from the Yanomami/Venezuela and other groups.

**Conclusions:** Our study revealed biomarkers, taxonomic and functional differences between the gut microbiome of Yanomami/Brazil and Yanomami/Venezuela individuals. This intra-Yanomami group diversity was accessed due to the increase number of individuals and group studied. These differences may reflect their semi-nomadic behavior, as well as, the local and seasonal diversity of the vast rainforest niche they explore, despite their shared cultural and genetic background. Overall, their microbiome profiles are shared with South American and African traditional groups, probably due to their lifestyle. The unique features identified within the Yanomami highlight the bias imposed by underrepresented sampling, and factors such as variations over space and time (seasonality) that impact, mainly, the hunter-gatherers. Therefore, to reach knowledge about human microbiome variations and their implications in human health, it is essential to enlarge data concerning the number of individuals, as well as the groups representing different lifestyles.

## Background

The transition of the traditional modes of subsistence to the current western lifestyles that occurred with the advent of modern practices (urbanization and industrialization) brought wide differences in diet and environment, factors proposed to be the main determinants of the gut microbiome composition. In fact, cross-population studies have demonstrated distinct taxonomic and functional profiles between the gut microbiome of hunter-gatherers/rural agriculturalists and urban-industrialized human groups. The main differences among the gut microbiome of westernized and non-westernized groups are that individuals from traditional communities harbor a more diverse gut microbiome, with higher levels of fiber-degrading bacteria, and unique taxa that are depleted in urban-industrialized populations [1–10]. The lifestyle aspects that characterize most of the westernized groups, concerning diet, environment, sedentary practices, among others, shape the gut microbiome, defining some taxonomic profiles that have been associated with an increased risk of metabolic and chronic disorders that affect modern populations [11]. Studies focusing on the differences between traditional and urban-industrialized groups may reveal diets and bacterial/archaeal taxa that can be helpful in the development of prebiotics and probiotics for modern disorders prevention and treatment. Considering that human groups that live in a non-western lifestyle are in decline, the study of the remaining traditional groups constitutes an extraordinary opportunity to explore and unravel the human gut microbiome before modernization.

The Amazon region is the largest tropical wilderness area in the world, covering ~ 7 million km^2^. This region includes the most extensive and preserved rainforest in the world (the Amazon Rainforest), vast areas of scrub-savannah that dominate the headwaters of the Brazilian and Guyana shields, as well as the Andes highlands, which are characterized by tundra-like grassy tussocks called the Puna [12]. Moreover, this region presents elevations ranging from sea level at the river’s mouth to an altitude of 6,500 meters in the Andes [13]. Having such high variable geomorphology, climate and vegetation cover, and harboring estimated 400-500 indigenous Amazonian Indian groups (Amerindians), this region offers a unique scenario for microbiome studies. These groups live in the same geographical region but explore distinct ecological niches, present distinct dietary habits, culture, language and degrees of isolation [14, 15]. This diversity may reflect in the gut microbiome composition and functionality, expressing the adaptation to evolutionary and ecological constraints of each site inhabited, despite being non-western populations.

The Yanomami is the largest indigenous semi-isolated group in the Amazon to maintain a traditional system of production based on hunting, fishing, gathering, and swidden horticulture [16]. They inhabit an area of 192,000 km^2^ in the Amazon region encompassing the Brazil and Venezuela border. Of the estimated 40,000 Yanomami, approximately 26,000 live in the 37,260 m^2^ reserve in Brazil and another 16,000 in Venezuela. Even though they represent a single semi-nomadic ethnic group of hunter-gatherers, they speak four different languages of the same family and live in villages located at sea level as well as on high mountains in a huge area in the Amazon [17]. Their diet is low in fat and salt, and high in fruits, fiber, and sylvatic animals. Atherosclerosis and obesity are virtually unknown among semi-isolated Yanomami, having low blood pressure, with no apparent increase as they age [18].

A previous study with uncontacted Yanomami from Venezuela revealed some aspects from the gut microbiome of this group [4]. In order to go deeper into the characterization of gut microbiome among traditional subsistence groups, we studied semi-isolated hunter-gatherers Yanomami individuals from Brazil. For this, we generated and analyzed metagenomic data of Yanomami from Brazil, and performed comparative analyses with those from Venezuela, other traditional groups from the Amazon (the Matses and the Tunapuco), as well as an urban-industrialized group (Figure 1). The Matses and the Tunapuco inhabit the borders of Peru that comprise the Amazon Region, but their lifestyles and environment are strikingly different. The Matses are traditional hunter-gatherers living by the sea level, while the Tunapuco is a rural agriculturalist community situated in the Andes highlands [6].

**Figure 1:**
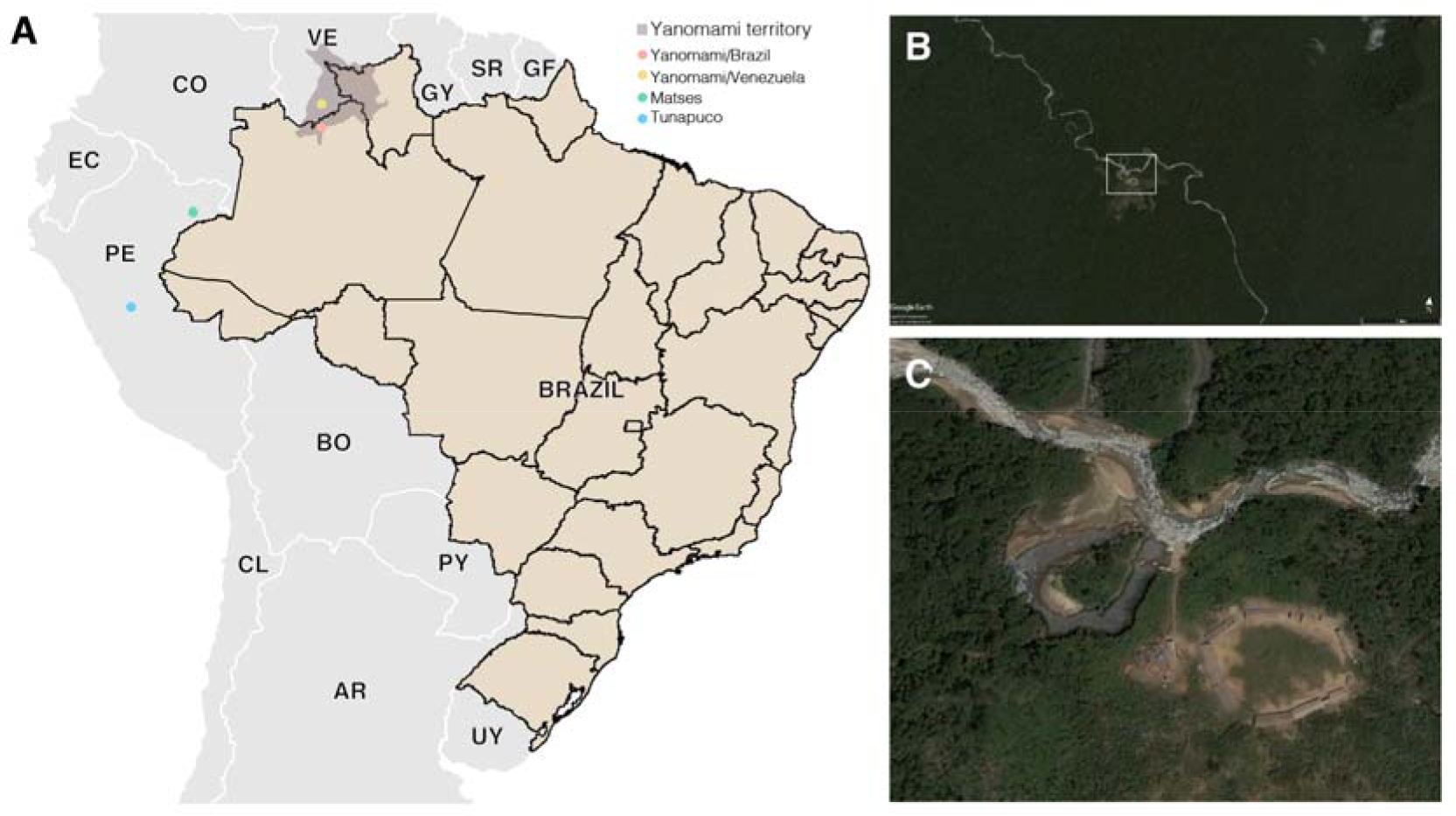
(A) Geographic locations of the South American traditional groups (B and C) Satellite image of a Yanomami village in the Brazilian Amazon. Source: Google Earth and Instituto Socioambiental (https://acervo.socioambiental.org/)

We hypothesized that, since the previous Yanomami group studied was uncontacted and lived in a remote area in Venezuela [4], their gut microbiome would present unique features in comparison with the large semi-isolated Yanomami group living in the Brazilian Amazon. Even though they are hunter-gatherers, their diet varies depending on the niche explored, since they live in areas ranging from near rivers to frankly mountainous regions. Moreover, we explored the taxonomy and functionality of these microbiomes, contributing to the understanding of features that can affect the health outcomes observed in modern populations.

## Results

### Intra and Inter Individual Diversity of the Gut Microbiomes

To unravel the gut microbiome diversity of Yanomami from Brazil individuals (Yanomami/Brazil, n = 15), we performed alpha and beta diversity analyses based on the bacterial genera profile identified by Kraken. For these analyses, we also reanalyzed and compared gut microbiome data gathered from other South American traditional communities: the Yanomami from the Venezuelan Amazon (Yanomami/Venezuela, n = 8) [4], the Matses from the Peruvian Amazon (n= 24) [6], the Tunapuco from the Andean highlands (n= 12) [6]; and a representative group of urban individuals from United States (US, n= 44) [19,20].

There was no statistically difference between the Yanomami/Brazil and Yanomami/Venezuela regarding intra and inter diversity (alpha- and beta-diversity, respectively) of gut microbiome, although the group from Brazil had the lowest alpha-diversity values. However, the Yanomami individuals showed the lowest bacterial alpha-diversity among the traditional groups and all the traditional human groups presented higher bacterial diversity compared to the urban individuals (Figure 2a). Regarding the beta-diversity, the Yanomami/Brazil presented the highest interpersonal variation, and the urbans presented the lowest, although there was no significant difference with the Yanomami/Venezuela group (Figure 2b). A clear segregation was observed among the semi-isolated and westernized individuals (PERMANOVA, P=0.001) based on Principal Coordinate Analysis (PcoA) generated with Bray-Curtis distances. In addition, a higher dispersion of Yanomami/Brazil and Yanomami/Venezuela samples was observed, stressing their higher interpersonal variation (Figure 2c).

**Figure 2:**
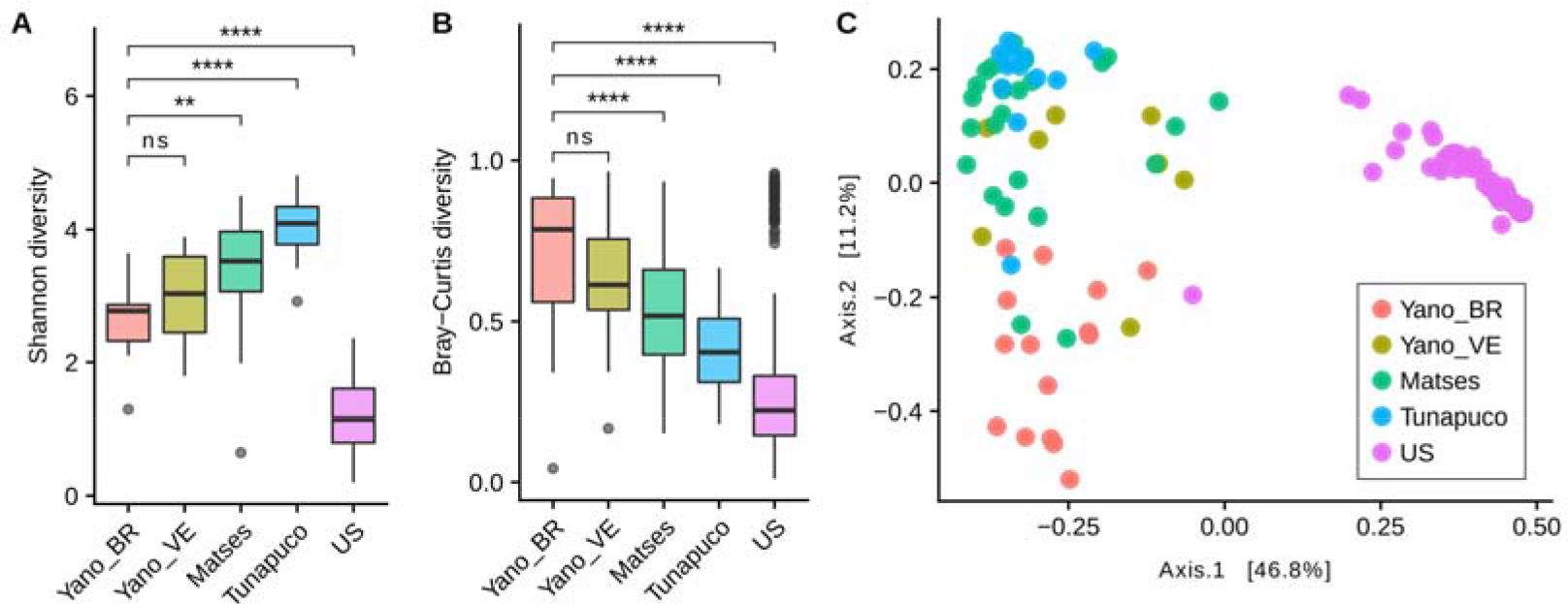
Alpha and beta-diversity comparisions of the gut microbiomes of each group. Analyses were perfomed on genus-level taxa tables. ns = not significant, **P<0.01, ****P<0.0001 (Wilcoxon test) (A) Boxplot of the Shannon diversity of each group (B) Bray-Curtis distances within each group (C) Principal coordinate analysis of Bray-Curtis distances. The colors of the boxplots and dots represent the different groups analyzed according to the legend. Yano_BR, Yanomami/Brazil; Yano_VE, Yanomami/Venezuela; US, US individuals.

### Microbiomes Taxonomic Characterization

In order identify which bacterial and archaeal taxa differentiate the traditional groups from the urban group, the microbiomes were compared at both phylum and genus scales. Thirty-two bacterial phyla were identified, with 16 phyla having significant differences in the relative abundances among the groups (Kruskal-Wallis test: P < 0.0001). Considering the traditional and urban groups, a clear difference at the phylum level was observed, with the former having a higher biodiversity characterized by *Firmicutes*, *Proteobacteria*, *Bacteroidetes* and *Spirochaetes*, while the urban group is mainly characterized by *Bacteroidetes* (Figure 3a).

**Figure 3:**
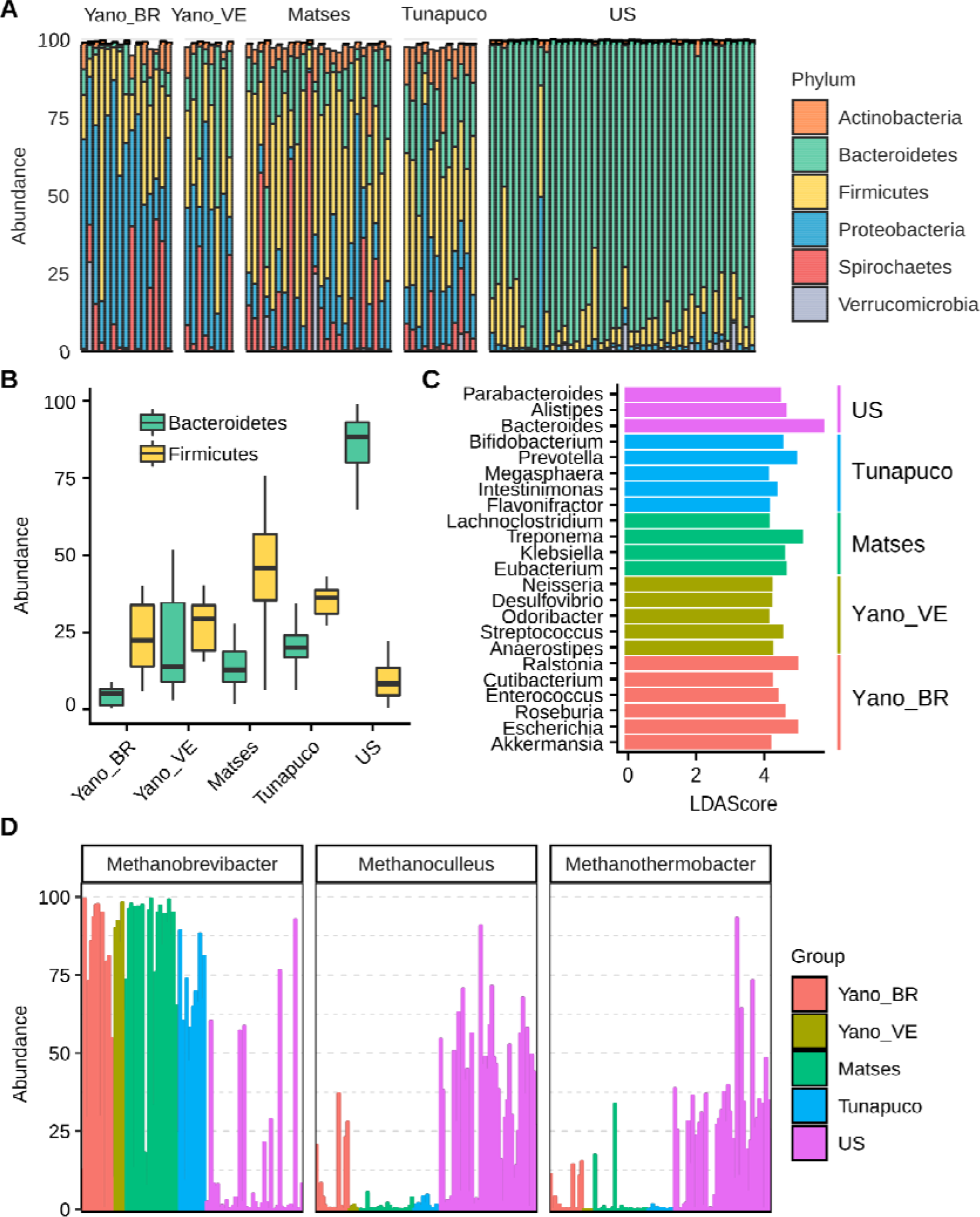
Bacterial and archaeal taxa differences among traditional and urban groups. (A) Barplot representing the relative abundance (percentage) of the most frequent phyla (B) Boxplots showing the *Bacteroidetes* and *Firmicutes* abundance (percentage) in each group (C) Bar chart showing the LDA scores > 4 of bacterial genera found to be significantly associated with each group (D) Relative abundance of the most prevalent archaeas identified in the groups. Yano_BR, Yanomami/Brazil; Yano_VE, Yanomami/Venezuela; US, US individuals.

The Yanomami/Brazil as well as the other traditionalists individuals follow a trend in which they have higher *Firmicutes* and lower *Bacteroidetes* levels, while the opposite was observed in the urban individuals (Figure 3b). Even though, the *Firmicutes* in each traditional group was characterized by distinct genera, no prevalent genus was consistently observed in the groups. In fact, all traditional groups presented different genera from the *Firmicutes* phylum as biomarkers (Figure 3c). Genera from *Bacteroidetes* phylum were demonstrated to be the biomarkers of the urban group (Figure 3c).

Distinctly from the other groups, *Proteobacteria* was the most prevalent phylum among the Yanomami individuals, despite their geographic origin (Brazil and Venezuela). The most abundant genera of this phylum in the traditional groups were *Escherichia* and *Klebsiella*, however, there is a contrasting higher abundance of *Escherichia* and *Ralstonia* genera in the Yanomami/Brazil, and therefore, they were defined as Yanomami/Brazil biomarkers. On the other hand, *Neisseria* and *Desulfovibrio* were defined as Yanomami/Venezuela biomarkers, while *Klebsiella* was the biomarker of the Matses group. It is noteworthy that *Cutibacterium* from the Actinobacteria phylum and *Akkermansia* from the Verrucomicrobia phylum were also deemed as the biomarkers of the Yanomami/Brazil (Figure 3c). Besides that, the Yanomami/Brazil, similarly with the other semi-isolated, present *Treponema* and *Brachyspira*, two genera from the Spirochaetes that were not detected in the urban group.

With respect to Archaea, we observed that the most abundant genus in the traditional groups was *Methanobrevibacter*, comprising ~70% of all archaea classified reads, while in the urban population, there were a high abundance of *Methanoculleus* and *Methanothermobacter*, all methane-producers archaea (Figure 3d).

### Microbiomes Functional Characterization

For functional characterization, the metagenomic reads of all groups were assigned to gene families from the SEED database, and were categorized for their functional roles in subsystems with 3 levels of resolution, in which level 1 represent the broader category. We observed a segregation between the traditional and urban groups regarding the abundance of functions at the level 3 subsystems. Interestingly, among the traditional groups, the Matses and Yanomami/Brazil individuals exhibits a clear segregation, however the Yanomami/Brazil present a more disperse pattern, indicating a quite diverse functional characteristic concerning functions at the level 3 subsystem (Figure 4a).

**Figure 4:**
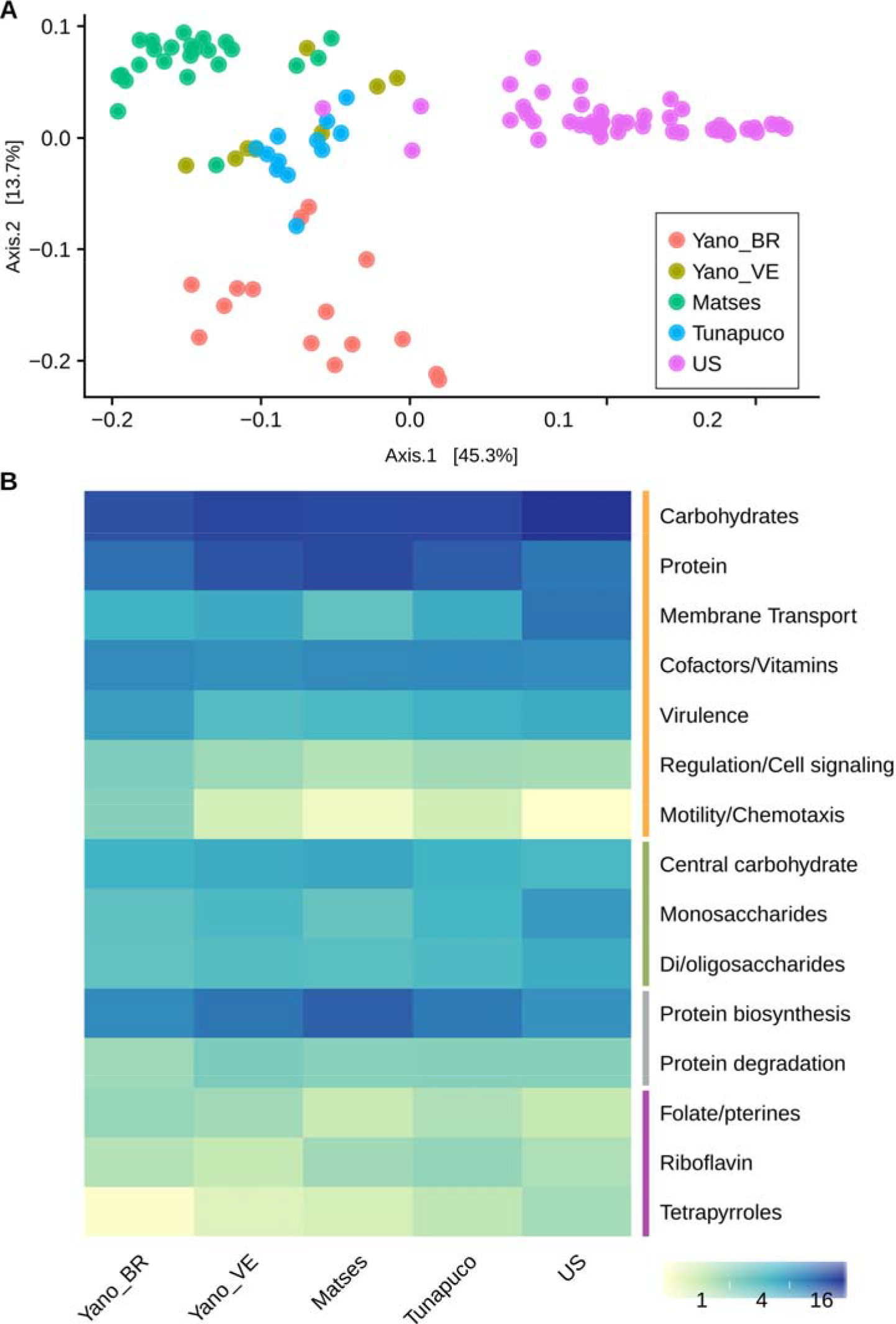
Functional metabolic characteristics of the traditional and urbanized microbiomes. (A) Principal coordinate analysis of Bray-Curtis distances based on functions at the level 3 subsystems. (B) Heatmap showing the main functions at level 1 (orange bar) and level 2 regarding carbohydrates metabolism (green bar), protein metabolism (gray bar) and cofactors, vitamins, prosthetic groups and pigments (purple bar). Yano_BR, Yanomami/Brazil; Yano_VE, Yanomami/Venezuela; US, US individuals.

The most abundant metabolic functions present in the microbiome of all groups at the level 1 were the metabolism of carbohydrates and proteins, while the membrane transport was also a main function in the urban group, but was depleted in the traditional groups (Figure 4b). We observed differences in the carbohydrate metabolic functions between the traditional and the urban groups. At level 2, the main carbohydrate metabolic functions in the traditional groups belonged to the central carbohydrate metabolism, with functions as pyruvate metabolism and glycolysis/gluconeogenesis being the most predominant; while in the urbans, the major functions were the monosaccharides and di-/oligossacharides metabolism (Figure 4b). In monosaccharides metabolism, there was also differences between the groups: D-galacturonate/D-glucuronate and xylose use were the most abundant functions in the traditional groups, while mannose metabolism was the most abundant in the urban group. The di-/oligosaccharides microbiome metabolism of US individuals is mainly driven by functions associated with lactose utilization, which was depleted in the microbiome of traditional groups. Regarding protein metabolism, there was no difference among the groups’ microbiome, with the most abundant functions being those associated with protein biosynthesis and degradation. We also observed differences in metabolic pathways related to cofactors, vitamins, prosthetic groups and pigments among the groups, the major gene families found in the microbiome of the Yanomami/Brazil and Yanomami/Venezuela were associated with folate/pterines, in the Matses and Tunapuco was riboflavin, and in the US group was tetrapyrroles (Figure 4b).

Interestingly, at level 1 subsystems, the microbiome of the Yanomami/Brazil was distinct from the other groups due to its significant higher abundance of gene families involved in regulation/cell signaling, motility/chemotaxis, and virulence (Figure 4b). The regulation/cell signaling functions in the Yanomami/Brazil is driven by abundance of programmed cell death and toxin-antitoxin systems, while the motility/chemotaxis function is driven by the presence of genes involved in flagellar motility in prokaryotes. The most abundant virulence function at subsystems 3 in the Yanomami/Brazil is cobalt, zinc and cadmium resistance.

## Discussion

The gut microbiome is a diverse ecosystem with multiple metabolic and immune functions associated with the diet and lifestyle of the host [11,21]. Therefore, considering the current variety of lifestyles and diets in human society, many aspects concerning the gut microbiome composition and functionality are yet to be accessed and explored to understand the influence of these factors in the gut microbiome. The study of human groups that still maintain traditional modes of subsistence (hunter-gatherers and rural agriculturalists) provides valuable information regarding the ancestral microbiome that existed before the urbanization and industrialization impacted human diet and lifestyle. So far, few studies explored worldwide traditional groups gut microbiome, and it is essential to enlarge data concerning the number of individuals, as well as different groups.

Therefore, in the present study, we characterized the gut microbiome of 15 semi-isolated Yanomami individuals from Brazil, and compared with other South American traditional groups (uncontacted Yanomami from Venezuela, the Matses and the Tunapuco) as well as an urban-industrialized group (US) [4,6,19,20], enlarging the number of Yanomami individuals analyzed, as well as of hunter-gatherers. These traditional groups explore different Amazonian niches and contrast with the US group, which lives in a densely populated urbanized and industrialized society with access to medical care and high hygiene standards.

Consistent with previous studies [1–10, 22], our analysis point to a higher bacterial diversity in the traditional groups, with diverse taxonomic and functional features that distinguish them from urban-industrialized individuals. Microbiomes harboring a high diversity showed a positive association with health, as consequence of the presence of a higher global metabolic potential, providing the host with a wide range of health-relevant metabolites [23, 24]. Besides that, the microbiomes of the traditional South American groups share features with traditional African groups (Western, Central and Eastern Africa): they are also more diverse than the urbans, are enriched in *Proteobacteria*, with the presence of some *Spirochaetes* that are depleted in industrialized populations (*Treponema* and *Brachyspira*) [1–10, 22]. Despite the South American and the African groups being in different continents and having distinct genetic origin, they maintain a traditional mode of subsistence and do not have access to processed and refined food in their daily diet [21]. This corroborates that population lifestyle and diet are the major determinants of the gut microbiome composition and diversity, overruling genetic backgrounds and geographic origin.

The taxonomic analysis of the South-American traditional groups demonstrated a common profile at bacterial genera level, even though each group presented a specific set of biomarkers. Interestingly, some biomarkers converge in their functional profile, e.g. *Roseburia*, *Anaerostipes*, *Eubacterium*, *Flavonifactor*, biomarkers of the Yanomami/Brazil, Yanomami/Venezuela, Matses and Tunapuco, respectively, are butyrate-producing bacteria. Butyrate is an anti-inflammatory short chain fatty acid (SCFA) that induces mucin synthesis, contributing to colon health and gut integrity [25, 26]. In contrast, the urban-industrialized biomarkers produce SCFAs other than butyrate, such as propionate, acetate, and succinate, which, in high proportions, may increase gut permeability, leading to a further unhealthy status [27]. Other biomarker of the Yanomami/Brazil group is Akkermansia, a mucin degrader, which has been associated with healthier metabolic status and better clinical outcomes [28,29].

Broader levels of functional categories poorly discriminated traditional and urban-industrialized groups. Interestingly, the stratification of these categories clearly segregated those groups. Differences were identified at level 3 of monosaccharides metabolism, where the main functions in the traditionalists and urbans were xylose and mannose metabolism, respectively. Xylans and Mannans, polysaccharides of xylose and mannose, are the two major classes of hemicelluloses that accumulate in plant secondary walls [30]. Interestingly, recent studies with mouse models revealed that mannose increased the Bacteroidetes to Firmicutes ratio in the gut, a characteristic observed in urban-industrialized groups [31]. On the other hand, Treponema, a prevalent genus in traditional populations that consume polysaccharide-rich diets, is a key xylan-degrader [32]. Another difference observed in the US versus traditional groups was the lactose utilization, which is enriched in the former and depleted in the latter group. This difference may be related to the lack of intake of dairy in the traditional groups [6]. Within the traditional groups, there was differences at level 3 of biosynthesis of vitamins: both Yanomami groups presented an enrichment in folate biosynthesis while the Matses and Tunapuco presented an enrichment in riboflavin biosynthesis, as well as in the US group. Riboflavin is the most commonly synthesized vitamin in the gut [33], and has been associated with the immune response through the activation of T-cells [34]. Folate is associated with high-fiber and low-fat diets [35], which agrees with Yanomami diet from the present study.

The microbiome of the Yanomami/Brazil is unique concerning the presence of higher levels of functions associated with virulence, driven by the cobalt, zinc and cadmium resistance. Cobalt is commonly distributed in nature and has a biologically role as metal constituent of the vitamin B12, however, excessive exposure induces adverse health effects [36]. Zinc is an essential nutrient and play a role in gene expression, biomolecular activity and structural DNA stabilization [37]. Cadmium is a non-essential element, representing an environmental hazard to human health when contaminates the food chain, causing cumulative toxic effects in diverse human organs [38]. Cadmium and Zinc are present in mine discharges, which disperses into air, water and soils, contaminating areas nearby mines [39]. However, in Yanomami/Brazil area, Cadmium contamination may occur as consequence of the continuous discharge of batteries anywhere by the Yanomami along decades.

## Conclusions

Exploring the gut microbiome of traditional groups is challenging, mainly due the difficult to access them. These groups are important, since they represent living representatives of ancestral behaviors/dietary long lost for a long time in the westernized groups. Our study revealed that even within very close and related traditional groups (as Yanomami/Brazil and Yanomami/Venezuela), there are taxonomic differences that distinguish their gut microbiome. These variations may reflect their nomadic behavior, as well as, the local and seasonal diversity of the vast rainforest niche they explore, despite their shared cultural and genetic background. Overall, their microbiome profiles are shared with South American and African traditional groups, probably due to their diet and lifestyle. This highlight the need to characterize larger sampling of human microbiomes, considering not only distinct lifestyle but also a broad population representing a particular lifestyle. Thus, we expect novel insights into the diverse factors that are associated with microbiome composition and human health.

## Methods

### Study Participants and Sample Collection

The protocol of this study was reviewed and approved by Oswaldo Cruz Foundation’s Ethics Research Committee N º 638/11 and by the National Ethics the Committee in Research – CONEP Nº 16907. Before participating in the study, a bilingual interpreter (a Yanomami native who spoke Portuguese) explained the leaders and/or Indigenous representatives, the purpose and importance of the study, the procedures to be carried out and finally requested permission by fingerprint consent of each participant. Participants were requested to provide a morning faecal sample and a labelled screw-capped plastic container was provided. A single stool sample was collected from each subject on the following day and samples were stored in separate sterile feces containers. At the time of the collection, age and sex information of the individuals were also acquired. These details are summarized in Table S1.

### DNA extraction, library preparation, and sequencing

Total DNA was extracted from 15 stool samples with FastDNA^®^ SPIN Kit (MP Biomedicals), following the manufacturer’s instructions. DNA concentration were evaluated using Qubit^®^ 2.0 Fluorometer (Life Technologies). Metagenomic libraries were constructed with TruSeq DNA Sample Preparation v2 Kit following the standard protocols. Purified libraries were sequenced on a HiSeq^®^ 2500 sequencer (Oswaldo Cruz Foundation High-throughput sequencing Platform) in two batches, producing a total of ~ 219 million reads, with an average of ~ 14 million reads per sample.

### Bioinfomatic Processing

Raw reads were trimmed and filtered (phred quality < 20, length < 30) using Trimmomatic [40]. The remaining reads (~ 206 million reads) were mapped to a human reference genome (Hg38) using Bowtie2 [41]. Non-host reads (~ 198 million reads) were used in further analysis. Besides the metagenomes generated in this study, we also analyzed shotgun metagenomic data from previously published studies: two hunter-gatherer communities (Yanomami from Venezuela, n=8 [4]; Matses, n=24 [6]), a rural agricultural community (Tunapuco, n=12 [6]) and urban populations (USA, n=44 [19,20]). These datasets were sequenced on Illumina plataforms and bioinformatic processing was performed in parallel with the data generated in this study.

Taxonomic classification was performed by Kraken [42], using a database of whole genomes of bacteria and archaea from NCBI. Functional classification were classified by SUPER-FOCUS [43] based on the genes families from the SEED database. Linear discriminant analysis (LDAs) were performed using LEfSe [44] to detect bacterial genera that characterize the differences between the groups (LDA score of > 4.0).

For general data manipulation and statistical analysis we employed the vegan [45] and phyloseq [46] packages in R. Shannon index of alpha-diversity was estimated for each metagenome, with pairwise Wilcoxon test being used for statistical difference evaluation. Beta diversity was estimated using Bray– Curtis dissimilarity and permutational multivariate analysis of variance (PERMANOVA) were performed with 999 permutations to estimate a P-value for differences among traditional and westernized groups.

## List of abbreviations

LDA: Linear discriminant analysis
LefSe: Linear discriminant analysis effect size
PcoA: Principal coordinates analysis
PERMANOVA: Permutational multivariate analysis of variance
SCFA: Short chain fatty acid

## Declarations

### Ethics approval and consent to participate

The protocol of this study was reviewed and approved by Oswaldo Cruz Foundation’s Ethics Research Committee N º 638/11 and by the National Ethics the Committee in Research – CONEP Nº 16907.

### Consent for publication

Not applicable.

### Availability of data and materials

The quality-filtered metagenomic sequences are available on the NCBI under the BioProject PRJNA527208.

### Authors’ contributions

JO collected the samples. LCC processed the samples and analyzed the data. LCC and ACPV interpreted the data and drafted the manuscript. All authors revised it. All authors approve the final version to be published and agree to be accountable for the work.

## Acknowledgements

We are especially grateful to the Yanomami people and we also thank the health personnel of the Distrito Sanitário Especial Indígena Yanomami for overall support during field work. We are particularly thankful to Dr. Edson Delatorre for the discussion of the manuscript and help with a figure.

